# Postnatal Development of the Gray Short-Tailed Opossum (*Monodelphis domestica*): Implications for Metatherian Decline at the K–Pg

**DOI:** 10.64898/2026.03.05.709990

**Authors:** Aidan M. C. Couzens, Clive Lau, Karen E. Sears

## Abstract

Marsupials give birth to extremely altricial offspring which must be reared externally for an extended period, often in a pouch. Despite often being considered a defining feature of marsupials, around a third of living species lack a pouch. Here, we describe the postnatal development of the gray short-tailed opossum, *Monodelphis domestica*, a small pouchless South American didelphid and consider its implications for life history evolution within Metatheria. We find that at birth, ossification and chondrogenesis in neonatal *M. domestica* is more extensive than in basal pouched Australidelphian marsupials like the dunnart. Key precocial milestones such as tarsal ossification, eye opening, growth of body fur, and chewing tooth eruption occur earlier and more rapidly. Principal component analysis of life history and reproductive traits reveals a pronounced r- to K-selected gradient across living marsupial species. Stochastic character-mapping based ancestral state reconstruction suggests that absence of the pouch, and by inference possession of an r-selected life history strategy characterised by large litters, short attachment phases, and accelerated weaning was likely ancestral amongst crown-group marsupials. The more K-selected reproductive strategy of pouched marsupials wherein there is a prolonged postnatal development window, and relative few young are produced, likely evolved during the early Cenozoic, and separately amongst australidelphian and ‘ameridelphian’ marsupials. Rather than making early marsupials more sensitive to environmental disturbances, we hypothesis that their possession of an ‘r-selected’ life history strategy may have been a key factor in their persistence through the K–Pg extinction.

## Introduction

Marsupials are one of the three major groups of living mammal, alongside the placentals and egg-laying monotremes (Luo, 2007). Compared with their sister therian group, the placentals, marsupials are unusual for giving birth to extremely altricial offspring after an abbreviated gestation period, sometimes as short as 10 days (Tyndale-Biscoe and Renfree, 1987). This is followed by an extensive postnatal interval until weaning, sometimes exceeding 6 months, during which time the young may be reared in a pouch (Tyndale-Biscoe, 2005). At birth marsupial neonates are effectively embryonic, comparable to 12-to-14-day old embryonic mouse (Hughes and Hall, 1988). In many taxa, the neonate skeleton is almost entirely cartilaginous, with a tube-like mouth, rudimentary lungs and gut, and paddle-like hind-limbs which are fused to the tail (Hill and Hill, 1955; Cook et al., 2021). Despite this extremely underdeveloped state, at birth marsupial neonates must attach to the mother’s nipples, sometimes by an extensive crawl, and then remain attached suckling for several weeks (Tyndale-Biscoe, 2005). Reared externally like this, marsupial neonates are vulnerable to external stressors such as hypothermia, desiccation, infection, and malnutrition (Smith and Keyte, 2020). To help shield the offspring, most living marsupials possess specialised abdominal folds which form a protective pouch; the ‘marsupium’ (Tyndale-Biscoe and Renfree, 1987; Edwards and Deakin, 2012). However, pouch structure varies widely across marsupials (Edwards and Deakin, 2012), and several American marsupial groups, such as shrew opossums and mouse opossums lack a pouch entirely or possess only a rudimentary seasonal one (Nowak 1999).

Today marsupials constitute only around 5% of living mammal species, with the rest, aside from a handful of monotreme taxa, belonging to the placental radiation (Burgin et al., 2018). In addition to being less taxonomically diverse than placentals, marsupials are also entirely restricted to the Americas and Australasia (Nowak, 1999). Despite being a relatively minor faunal component everywhere except Australasia today, during the late Cretaceous marsupials and their close relatives (Metatheria) were taxonomically and ecologically diverse, and widely distributed across North America, Asia and Europe (Archibald, 1983; Wilson, 2013; Williamson et al., 2014; Wilson et al., 2016). Following the Cretaceous–Paleogene mass extinction the diversity of metatherians declined markedly, and they were at least partly replaced by placental mammals (Lillegraven and Eberle, 1999; Wilson, 2013; Williamson et al., 2014). The unusual mode of marsupial development and life history has been cited as one factor in this broadscale decline, with factors such as their reliance on extensive external rearing being hypothesised to have increased their vulnerability to environmental change during the end-Cretaceous (Lillegraven et al. 1987). It has also been hypothesised that due to their shortened gestation period, extended oral attachment, and early specialisation of the forelimbs, early marsupials may have been adaptively constrained compared with placental counterparts (Lillegraven, 1975; Lillegraven et al. 1987; Sears, 2004; Goswami et al., 2016).

Here, we describe the postnatal development of laboratory reared individuals of the gray short-tailed opossum, *Monodelphis domestica*, a small (∼100g), pouchless marsupial native to central South America. Although, *M. domestica* is widely used in biomedical research across a range of areas including spinal cord regeneration (Wheaton et al., 2011), melanoma induction (Nair et al., 2014), teratogenesis (Molineaux et al., 2015), and was the first marsupial to have its genome sequenced (Mikkelsen et al., 2007), its postnatal development has not been described comprehensively. Due to its peculiarity as pouchless marsupial, *M. domestica* also offers and interesting comparison with better characterised pouched Australian marsupials like the tammar wallaby (*Macropus eugenii*) and the fat-tailed dunnart (*Sminthopsis crassicaudata*) (e.g., Renfree and Pask, 2011; Cook et al. 2021). Here we examine: (1) How does the absence of a pouch affect the rates and sequence of postnatal development in *M. domestica*? (2) When and why did pouchless life history strategies evolve? (3) What are the implications of this for our understanding of the role of life history and reproductive strategies in the evolution of marsupials and their close relatives? Our results suggest that possession of a pouch very strongly influences reproductive strategy, the timing of life history milestones, and rates of character development. Furthermore, lack of a pouch is recovered as ancestral for crown-group marsupials, suggesting that early marsupials lacked the very elongated external rearing, and low fecundity, often thought characteristic of marsupials. We consider implications of this revised view of marsupial life history for scenarios which have been proposed for the broader decline of metatherians since the end-Cretaceous.

## Materials and Methods

### Specimen Collection

Following mating, female opossums were monitored during the 14-day gestation period. The first 24 hours following birth was demarcated P0, 24–48 hrs as P1, and so on. Postnatal specimens were collected by gently removing attached young from the mother. For characterising gross anatomy and microCT scanning, individuals were euthanised using approved IACUC methods. Specimens were photographed in lateral view using a Nikon Coolpix P7700 digital camera with scale bar immediately following euthanisation. Specimens were collected daily at the same time from day 0 until day 10, and then at 5-day intervals until day 30, and 10-day intervals from day 30 until day 50. Photographs were colour balanced and contrast improved in photoshop. All animal husbandry and euthanisation activities were performed under IACUC approved protocol ARC-17-069 to KES at the University of California Los Angeles.

### Bone and Cartilage Staining

Following collection, specimens were placed in warm water for 30 seconds, following which skin and viscera were removed under a dissecting microscope. Specimens were then placed in ice cold 95% ethanol for 1 hour before being moved to fresh 95% ethanol and gently rocked over night at room temperature. Specimens were rinsed in 70% ethanol/ 5% acetic acid for 30 minutes and stained overnight in a freshly prepared stain solution. A stain solution of 0.4% alcian blue was created by dissolving 0.4g of alcian blue powder in 50% ETOH, swirling, and placing in a water bath at 37°C. After dissolution 25 mL of water was added and 65 ml of 95% ETOH. A second stock solution of 0.5% alizarin red S was created by dissolving 0.5g of alizarin red S in 100ml of water and swirling till dissolved. From these stock solutions the final stain solution was made freshly by combining 5 ml Alcian Blue stock solution with 5 ml glacial acetic acid, 70 ml 95% ETOH, 20 ml water. When ready for staining 100ul of Alizarin red stock solution was added to this 100ml volume. Observations were made under a Leica S9i dissecting microscope and camera.

### Ruthenium Red Staining and MicroCT Scanning

We followed the ruthenium red staining approach of Gabner et al. (2020) for contrast staining of bone and cartilage. Unlike embryonic mouse specimens, in opossum postnatals the skin needed to be removed for effective stain penetration. Specimens were soaked for 30 seconds in a warm water bath (∼80C), and skin was removed carefully under a dissecting microscope with forceps. Specimens were then fixed in 80% ethanol solution for 7 days at 4°C. Staining was ineffective with formalin fixed specimens. Following fixation, samples were washed twice for 2 hours in 50% ethanol solution at 25°C in an incubator with a rocker. Next, specimens were stained in 0.1% solution of ruthenium red made by dissolving the ruthenium red powder slowly in distilled water to create a 0.2% w/v solution, after which an equal volume of 100% ethanol was added slowly using a magnetic stirrer. Specimens were then stained at 25°C on a rocking table for at least 72 hours. For later postnatal specimens longer staining times up to 120 hours were required. Stain penetration was checked via incision. After staining, specimens were washed in 50% ethanol solution for 2 hours, and a further 24 hours with gentle rocking, followed by at least two washes in 40% ethanol for two hours each. If after these steps, the solution was still pigmented, additional 40% ethanol washes were used. Next specimens were mounted in 1.5% low melt agarose in 40% ethanol in a 50ml falcon tube for scanning. Specimens were scanned at the Molecular Biology Imaging Centre within the Department of Radiology at the University of Southern California using a GE Phoenix Nanotom M scanner. Scans were acquired at voltage of 80kv, current of 120ua, and an exposure time of 750ms using a 0.01mm copper filter. Following reconstruction, image stacks were analysed with Avizo software. Volumes were cropped and voxels assigned to ‘bone’, ‘cartilage’, ‘teeth’, and ‘air’. Surfaces for each label were reconstructed using the ‘generate surface’ function and visualised with the ‘view surface’ function.

### Principal Component Analysis

Trait data related to reproduction and life history were collated for 89 marsupial species from Nowak (1999). Trait data included body mass(g), gestation length (days), weaning age (days), pouch structure (‘absent’, ‘folds’, ‘pouch’), litter size, timing of eye-opening, and breeding mode (‘continuous’, ‘seasonal’). Where data was given as a range, or for male and females separately, the median was computed. Using this data, we performed a principal component analysis in R. Due to large amounts of missing data for many of these traits, analysis was subsequently restricted to gestation length, litter size, weaning age, and pouch structure. Prior to analysis with the function ‘prcomp’ in the package “stats” we removed species with missing data and scaled the data using the argument ‘scaled=T’.

### Ancestral State Reconstruction

To reconstruct pouch evolution, we performed ancestral state reconstruction using stochastic character mapping technique. Three states were scored: absence of a pouch (‘A’), presence of a pouch (‘P’), or presence of abdominal folds (‘F’). From our sample of 89 marsupial species, all congeneric species shared the same pouch states, so we mapped generic level pouch states across 226 marsupial species present on the time-scaled DNA maximum clade credibility tree of (Upham et al., 2019). In addition, 16 placental and two monotreme taxa were included as outgroups, for a total of 244 phylogenetic tips. We used a stochastic character mapping approach (Bollback, 2006; Revell, 2013) implemented in R with code modified from Hughes et al. (2021). To account for variation in tree topology and divergence times, 500 stochastic character maps were estimated on a random sample of 100 trees drawn from the posterior tree distribution of Upham et al. (2019). This was implemented using the functions ‘simmap.parallel’ and ‘describe.simmap.alt’ developed by Hughes et al. (2021). The function ‘simmap.parallel’ takes a tree or a sample of trees, a discrete character matrix, a time-homogenous model of discrete character evolution (i.e., Q-matrix), and a root state prior. A ‘FitzJohn’ root state prior was assumed. Models of character evolution were selected by estimating a Q-matrix for the character matrix and the maximum credibility tree using the function ‘fitMk’ (Revell, 2012). Seven character transition models were assessed, including four custom models, and three generic models; all-rates different (ARD), equal rates (ER), and symmetrical rates (Table 1). The models range from more constrained to more evolvable models and differ in the types of transition between the three states, ‘no pouch’ (‘A’), ‘folds’ (‘F’) and ‘pouch’ (‘P’) they allow.

**Table 1.** Transition rates used for model selection and stochastic character mapping.

|  |  |  |  |
| --- | --- | --- | --- |
| A) M1- Two Rate (unconstrained symmetrical) |  |  |  |
|  | Absent | Folds | Pouch |
| Absent | . | 1 | 2 |
| Folds | 1 | . | 1 |
| Pouch | 2 | 1 | . |
| B) M2- One rate (constrained) |  |  |  |
|  | Absent | Folds | Pouch |
| Absent | . | 1 | . |
| Folds | 1 | . | 1 |
| Pouch | . | 1 | . |
| C) M3- two rate (constrained) |  |  |  |
|  | Absent | Folds | Pouch |
| Absent | . | 1 | . |
| Folds | 1 | . | 2 |
| Pouch | . | 2 | . |
| D) M4 – Four rate (constrained) |  |  |  |
|  | Absent | Folds | Pouch |
| Absent | . | 3 | . |
| Folds | 1 | . | 2 |
| Pouch | . | 4 | . |
| E) ARD - Six rate (all rates different) |  |  |  |
|  | Absent | Folds | Pouch |
| Absent | . | 1 | 2 |
| Folds | 3 | . | 4 |
| Pouch | 5 | 6 | . |
| F) ER – 1 rate (all rates the same) |  |  |  |
| Absent | . | 1 | 1 |
| Folds | 1 | . | 1 |
| Pouch | 1 | 1 | . |
| G) SYM – Three rates (forward and reverse rates the same) |  |  |  |
|  | Absent | Folds | Pouch |
| Absent | . | 1 | 2 |
| Folds | 1 | . | 3 |
| Pouch | 2 | 3 | . |

## Results

### Gross Morphology

At birth young are hairless, partly translucent, and approximately 10 mm in length (Fig. 1, Table 2). Some internal organs are externally visible, as is vasculature in the neck, head and abdominal region. The head is laterally flattened without a distinct neck. A pigmented retina is visible as are external openings for the oral cavity, ear, and nares. The region of the future ear pinnae is pale in colouration. The forelimbs are well developed, with a flexed elbow and grasping manus which can make powerful adduction motions. At birth, the manus has recurved claws on all digits. The hind limb is substantially smaller, paddle shaped and with only nub-like digits lacking claws. The thoracic, lumbar, and caudal region is strongly arched. The body is laterally narrow, and the young do not maintain a prostrate pose. The gut and internal organs appear simple. Dissection revealed heart, lungs, liver, stomach, pancreas, small and large intestine, and a small bladder. The tail is short and fused with the upper hindlimb.

**Table 2.** Major postnatal events in *Monodelphis domestica*. Abbreviations for tooth types: M=upper molar, m=lower molar, DP=Deciduous upper premolar, dp=deciduous lower premolar, I=upper incisor, i=lower incisor, C=upper canine.

| Postnatal day | Gross characteristics | Skeletal and dental characters | Behaviour |
| --- | --- | --- | --- |
| **P0** | Mouth tube-like and fused to mothers' nipple. Skin lacks hair and partly translucent. Pigmented retina. Well-developed fore-limb with flattened claws on distinct digits. Hind-limb fused to tail. Hind-foot paddle shaped with short, rounded digit precursors. Heart well developed but lungs flattened and small. Liver and stomach are large. Upper and lower intestine present but short. | Ossification in premaxilla, anterior maxilla, and anterior dentary. Incipient ossification in palate. | Crawl to nipples. Attachment and suckling. |
| **P1** | Neurocranium rounded. |  |  |
| **P4** | Hind-limb and tail start to separate. |  |  |
| **P5** | Lungs more vascularized. Hind-foot claws evident. |  |  |
| **P7** | Hair on head. | Alveoli for Mx/mx, C, post. lx and ix present. |  |
| **P8** | Distinct neck. Audible vocalisations. |  | Vocalisation |
| **P9** | Mouth starts to unzip anteriorly. |  |  |
| **P10** | Vibrissae on snout. Scrotum in males. Growth accelerates. |  |  |
| **P15** | Body hair. Ear pinnae separated. Mouth cleft extends past eye. Hindlimb increases in size relative to forelimb. Distinct hindfoot digits. | Enamel mineralisation well underway. Upper and lower molar crowns erupting. Anterior dentition including canines and incisors mineralising but unerupted. |  |
| **P20** | Mouth separated and mobile. Eye enlarged. Dorsoventral colouration. Opposable first digit on hindfoot. | Enamel mineralisation extensive. m2, m1, dp3, dp2, and lower incisors erupting, and M1 and DP1-3 erupting. Canines and upper incisors unerupted. |  |
| **P25** | Tail coiled. Snout elongating. |  | Pup detached from nipples. Maybe attached to mother's haunches with jaws and forelimbs. |
| **P35** |  |  | Eyes open. Pup attached to mother's haunches. |
| **P40** |  | M3/m3 and<br>M4/m4<br>incompletely<br>erupted |  |
| **P50** |  |  | Weaning |
| **P63** |  | Anterior dentition<br>completely<br>erupted. M3/m2<br>and M4/m4<br>unerupted. |  |

**Table 3.**
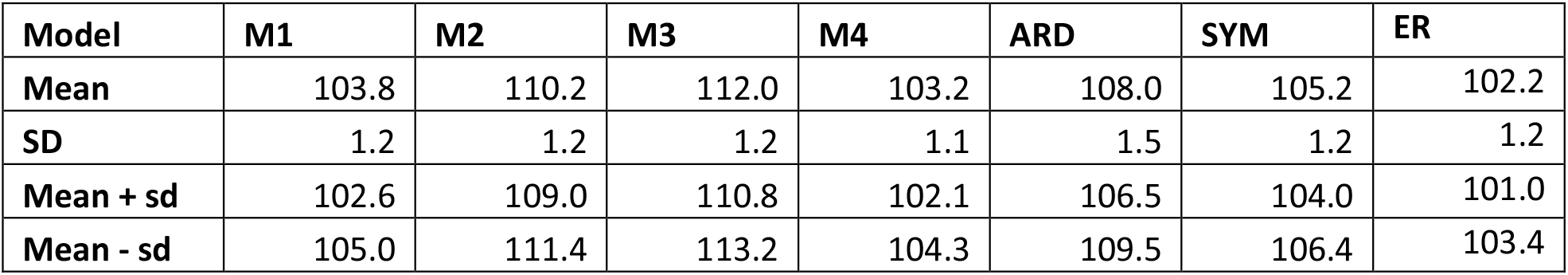
Akaike Information criterion scores for transition rate models fit with maximum-likelihood.

| Model | M1 | M2 | M3 | M4 | ARD | SYM | ER |
| --- | --- | --- | --- | --- | --- | --- | --- |
| Mean | 103.8 | 110.2 | 112.0 | 103.2 | 108.0 | 105.2 | 102.2 |
| SD | 1.2 | 1.2 | 1.2 | 1.1 | 1.5 | 1.2 | 1.2 |
| Mean + sd | 102.6 | 109.0 | 110.8 | 102.1 | 106.5 | 104.0 | 101.0 |
| Mean - sd | 105.0 | 111.4 | 113.2 | 104.3 | 109.5 | 106.4 | 103.4 |

**Figure 1.**
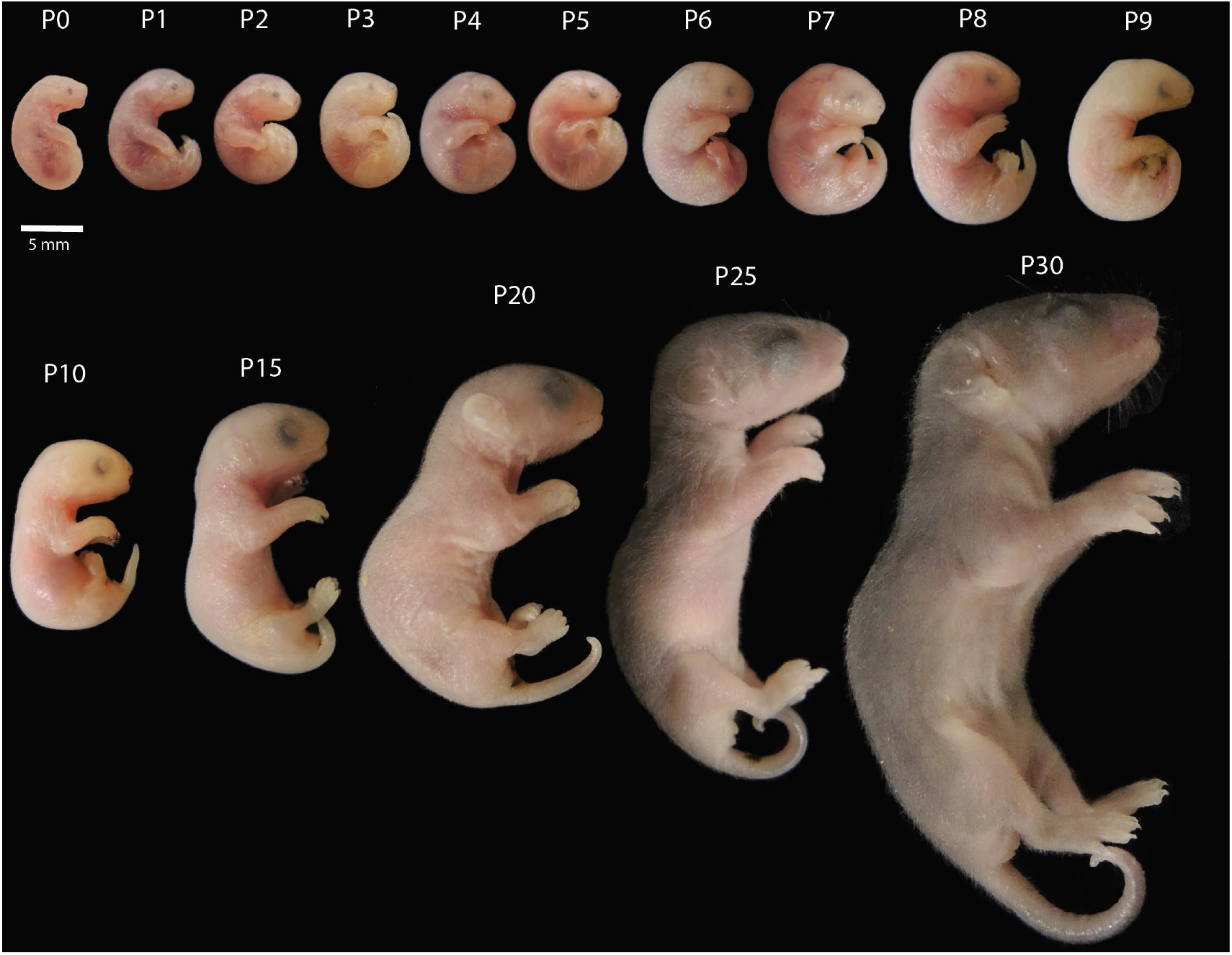
Gross morphology of postnatal *Monodelphis domestica* in lateral view extending from birth until 30 days post birth. Numbers are postnatal (P) days after birth with P0 being day of birth.

During the first five days of postnatal development there is little outward change in gross anatomy or size (Fig. 1). By P1, the neurocranium has a more domed appearance in profile. The liver is lobate and intestinal coils are present. The heart takes up much of the thorax and is similar in size to lobes of the lung. By P3, a distinct fold is present on the anterior border of the mouth, adjacent to the oral shield, marking the site of upper and lower jaw separation. The organs fill the abdominal cavity more extensively. By P4, the first digit on the forehand appears shorter than the other manual digits. The tail and upper hindlimb start to separate. Lungs are wedge shaped rather than flattened and have distinct alveoli. By P5 the cleft forming between the upper and lower jaw is deeper and extends more posteriorly. The nose has a darker pigmentation than the surrounding snout. The hindfoot digits exhibit evidence for nails. A genital protuberance can be observed anterior to the cloaca. The lungs have a darker colouration. By P6, there is noticeable increase in overall size compared with P5. A distinct thickening is present in the ear region. On the manus, pads are present and rounded elongate claws tip digits. The hindlimb and tail are separated. The internal organs are larger. At P7, a distinct fold is observed at the base of the nostril and an eye-lid cleft is visible. The first-digit on the hindfoot is clearly shorter than the other digits. Small hairs are visible on the dorsal rostrum. By P8, a distinct neck is apparent (Fig. 1). Pads are evident on the plantar surface of the hindfoot, although the digits are still connected by integument. Vocalisations are audible. At P9, the oral fold extends under the eye, and the mouth has started to separate anteriorly. The oral shield is reduced in size. Vibrissae are present around the eye and distinct rows for vibrissae are present either side of the rostrum. The forelimb digits are well separated but remain webbed on the hindfoot. Small, rounded nails are visible on the hindfoot digits.

At P10 small whiskers are present on the muzzle and hair extends to the dorsal part of the head (Fig. 1). Eyes are becoming enlarged. Hindfoot digits are becoming separate. The tail is straighter and more elongate and has a distinct tip. From P15 onwards, growth of the body accelerates, and the snout starts to elongate. At P15, hair is present on both the upper and lower jaw and on all body regions. Long whiskers evident on snout and lower jaw. The rounded ear pinnae are separated from the head. Digits are elongate on the hindlimb and there are well-developed claws. A defined oral cleft extends under the eye, but upper and lower jaw are not yet separated. In males, testes are sack-like with a distinctly darker colouration than surrounding belly. In females, what we interpret as two branches of the uterus are apparent. At P20, body length has increased markedly compared with P15 (Fig. 1). Eye lids are visible. The lower jaw is separated and mobile. Vibrissae on the snout and lower jaw are more elongate. Dorsal colouration is darker than ventral colouration. First digit on hindfoot shows evidence of opposability. Last third of tail is dorsoventrally flattened and the tail tip is recurved. At P25, pups can be observed detached from mother. The eye has become larger, and the snout is more elongate, creating a flatter nasal profile. The region below and behind the eye is swollen but lacks vibrissae. Fur cover is thicker and the contrast between dorsoventral colouration of the pelage is stronger. Unsheathing of hindlimb from the body wall creates distinct knee. Tail is coiled. At P30, body length has almost doubled from P25. The snout has elongated further, and the ear pinnae is more unfolded. Prominent skin folds have formed on the ventral surface of the body. Feet and hands are more flattened in profile. By P35, eyes are open, and pups were observed separated from the mothers’ nipples having relocated to her haunches. Forelimbs and jaws used to grasp fur. At P40 pups were clinging to the mothers’ sides or were independent amongst cage litter. By P50 pups are essentially adult-like in gross morphology, although they have not yet attained adult size.

### Postnatal Dental and Skeletal Development

At birth, ossification observed in the premaxilla, anterior maxilla, and anterior dentary (Fig. 2B). Faint alizarin staining present in the palate. The hyoid and tracheal cartilages are apparent. No teeth are mineralised. No clear evidence for extensive tooth bud initiation. Cartilaginous precursors for most postcranial elements are present (Fig. 2B). In the forelimb, ossification is incipient in the scapula, clavicle, humerus, radius, ulnar, and distal phalanges. The more proximal limb elements are better ossified. Remainder of the skeleton is unossified. In the hindfoot and distal tail some elements not chondrified. In the hindlimb, the tarsals and digital cartilages are present, but excepting the first digit, the hindfoot is not separated into distinct metatarsals and phalanges. Condensations for 5 tarsal elements are observed, but no evidence for a prehallucal condensation. In the pelvic region, condensations for the ilium, pubis, and ischium. Faint cartilaginous condensations for the epipubic bones. The cartilaginous vault of the braincase is open dorsally.

**Figure 2.**
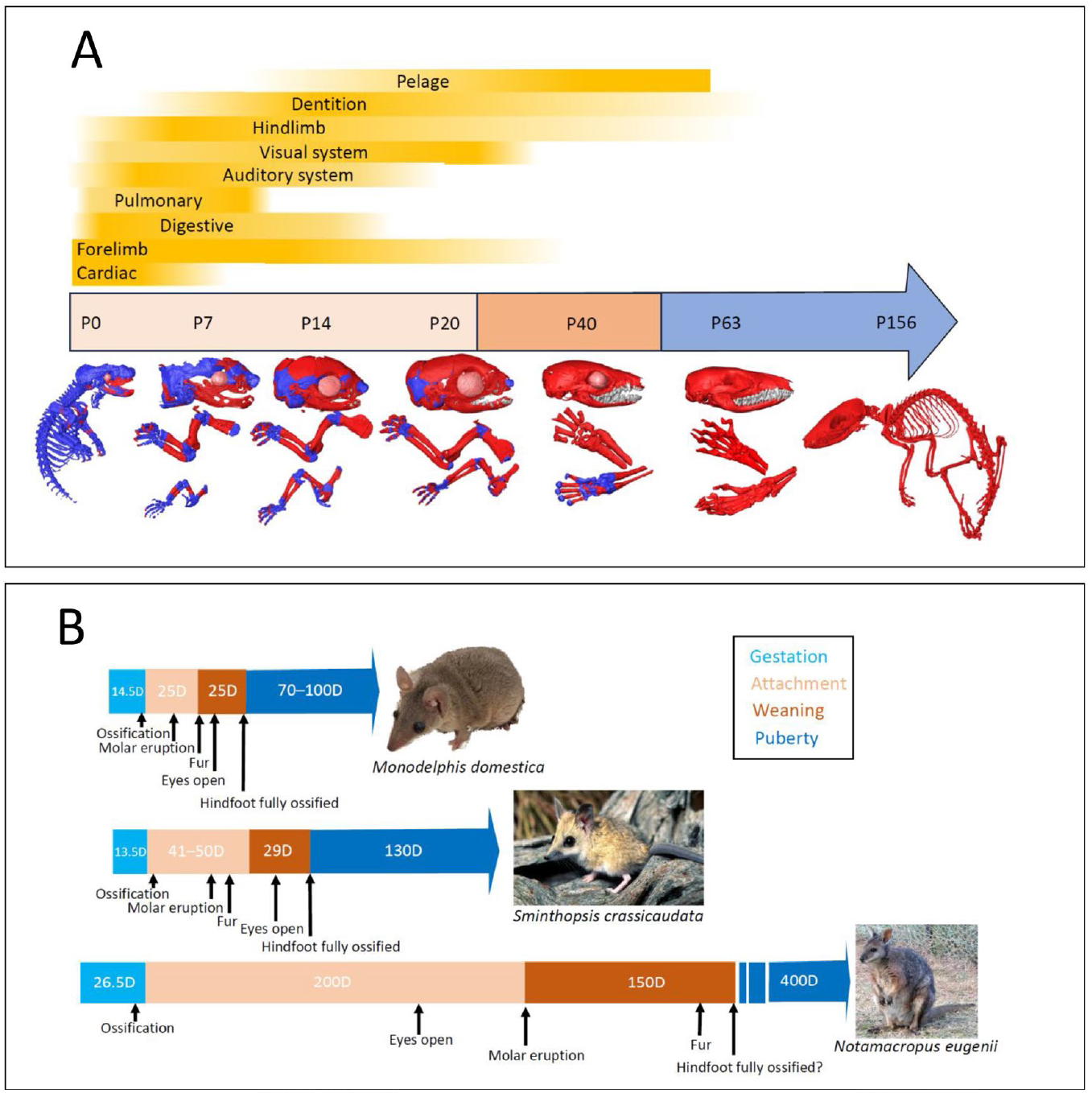
**A**. Relative stages of postnatal organ system and cartilage (dark blue) and bone (red) development. Light beige box = birth to detachment. Dark beige box = detachment to weaning. Blue arrow = weaning until adulthood. **B**. Comparison of the duration and timing of postnatal milestones in the grey short-tailed opossum (*Monodelphis domestica*), the dunnart (*Sminthopsis crassicaudata*), and tammar wallaby (*Notamacropus eugenii*).

By P1, ossification in the upper and lower jaw has extended posteriorly. Ossification is evident in the premaxilla, palate, exoccipital, basioccipital, ectotympanic, squamosal, and pterygoid. The clavicle, scapula, humerus, radius and ulnar are extensively ossified. Anterior regions of the vertebrae including all the cervical vertebrae (including axis and atlas) and thoracic vertebrae 1-to-13 show ossification. For the vertebrae, ossifications were well developed in the centrum and transverse processes. Vertebral ossifications are present before ossification in the ribs, and ossifications in the ribs initiate first proximally.

At P4, there was ossification in the frontal bone directly above the orbit (Fig. 2B). This ossification ends at the dorsal border of the cranial vault which remains open. Vacancies are observed in the dentary, maxilla, and premaxilla for developing teeth. The bone in these regions is thin and netted in appearance. By P5, the malleolus and ectotympanic are distinct. Large vacuoles in the dentary for the lower molar. Ossification is underway in the squamosal at the rear of the orbit. Ossification is present at the dorsal posterior margin of the nasal bone and in the lachrymal. Ossification centres in the vertebrae extend to the 5^th^ lumbar vertebrae. Cartilaginous elements in the hindfoot are distinct. All phalanges, metatarsal and tarsal elements are present including the prehallux. Anlagen for the caudal vertebrae are present. By P7 vacancies are present in the dentary and maxilla for tooth buds. In the maxilla and premaxilla, evidence for canine and incisor tooth buds.

At P10 ossification of the cranium much more extensive (Fig. 2B). In the forelimb, the metacarpals and phalanges are ossifying, although no ossification observed in the carpals. In the hind limb, the distal phalanges (claws) are ossified in digits 2, 3, and 4. In the pelvis, the pubis and ilium are substantially ossified. In the vertebral column, all the lumbar vertebrae show ossification as do caudal vertebrae 1–8. At P14, tooth mineralisation well underway. Upper and lower molar crowns erupting. Anterior dentition including canines and incisors mineralising but unerupted.

By P15 there are ossification centres in all the caudal vertebrae. In the hindlimb, ossification is underway in the metatarsals, the calcaneus, the central phalanges of digits 2–4, and the distal phalange of digit 1, and the prehallux. The other tarsal elements are present as distinct cartilaginous elements. In the forelimb, ossification is more advanced, with ossification present in all phalanges as well as the metacarpals. However, amongst the carpals ossification only observed in the pisiform. In the skull, the membranous bones of the skull are starting to ossify creating webs of thin bones in the frontal and parietal. In the pelvis, the ilium, pubis, ischium and epipubic bones are well ossified but still maintain thick epiphyses. The acetabulum is ossifying but the proximal and distal femur are cartilaginous.

At P20, first ossification is evident in the carpals and tarsals. In the carpals, ossification is observed in the prehallux, the proximal pisiform, and proximal edge of the hamate. Among the tarsals, a small ossification centre is evident in the centre of the calcaneus. The intercostal centra are ossified but have thick cartilaginous joints between them. A proximal fibular sesamoid is evident but not ossified. The caudal centra are ossifying and there are cartilaginous precursors for the spinous, transverse, and haemal processes. In the cranium, the maxillary bone is vacuous with large vacancies for prospective teeth but also spaces in front and slightly above the eye. The premaxilla and maxilla are not fused. Teeth are mineralised in the upper and lower jaw with no contact between the upper and lower dentition. Tooth mineralisation is extensive. On the lower jaw, m2, m1, dp3, dp2, and lower incisors erupting and on the upper jaw M1 and DP1-3 erupting. Canines and upper incisors unerupted.

By P25, most cartilage has been replaced by bone in the skull, but sutures remain open throughout, especially on the dorsal surface between the frontal and parietal and along the mid-line suture. Extensive cartilage still present in the lower jaw. In the hindlimb, the phalanges and metatarsals are mostly bone, but aside from the calcaneus all the tarsals are largely unossified. In the forelimb, the phalanges and metacarpals are well ossified, but there are ossification centers in several carpals including the trapezium, the hamate, the lunate, the pisiform, the capitate, and the scaphoid. In the hindfoot, alcian blue staining is notably concentrated along the prospective articular surfaces, especially between the tibia and astragalus, and in interphalangeal joints. At P40, effectively the entire anterior dentition has erupted and only the M3/m3 and M4/m4 incompletely erupted.

### Phylogenetic Analysis

Model fit comparisons across seven transition models revealed significant variation in model performance based on Akaike Information Criterion (AIC) scores (Table 2). Generally, there were low rates of variation in model fit despite differences in tree topology and branch lengths. Based on AIC ranking, the highest performing models were the equal rates (‘ER), M4, and M1 models. These models fall within a range of ±2 Δ AIC (102.2 – 103.8) indicating no significant differences in model performance. The lowest ranked model was M3 (112.1 ± 1.1 AIC) which was ± 10.1 Δ AIC below the best ranked model. The best ranked models were simpler, and more evolvable models of character evolution compared with the lowest ranked models. Complex, but highly evolvable models like ARD received intermediate rankings.

Ancestral state reconstructions using the best ranked models recovered consistent patterns of pouch evolution. Under ER, the ancestor of both Theria, and Marsupialia, is recovered as likely lacking a pouch. A pouch is reconstructed as evolving twice, once within Didelphimorphia, and once amongst Australidelphia, with caenolestids (Paucituberculata) retaining the ancestral therian condition of lacking a pouch. Amongst Didelphimorphia, likelihoods are ambiguous as to whether the pouch was ancestrally present amongst all members of the group and lost amongst mouse opossums + short-tailed opossums or only evolved amongst a clade comprising all other didelphids.

Compared with the best ranked models, lower ranked models recovered broadly similar patterns of character transition. For instance, the lowest ranked model, M3, still recovered strongest support for ancestral absence of the pouch amongst crown-group marsupials, and the Australidelphia-Didelphimorphia node. However, there was more ambiguity in these reconstructed states with stronger support for the presence of the pouch. The M3 models recovers much higher ambiguity in derived crown group nodes, with stronger support for reversals of pouch loss and gain amongst diprotodont marsupials. Amongst Didelphimorphia, loss of the pouch is inferred to involve transition through a step with folds.

## Discussion

Here, we have examined how absence of the pouch influences postnatal development with the grey short-tailed opossum, *Monodelphis domestica*. Our results show that absence of the pouch is associated with earlier initiation, and more accelerated appearance of key postnatal milestones relative to other marsupials (Fig. 2B). At birth, *M. domestica* exhibits more extensive cranial and postcranial ossification (Fig. 2A) compared with basal pouched marsupials like the dasyuromorphian fat-tailed dunnart, *Sminthopsis crassicaudata* (Cook et al., 2021). There was earlier evidence for the onset of cranial (maxilla, dentary, palatine) and postcranial (radius, humerus, and distal phalanges) ossification, compared with the dunnart. As with previous studies, the general sequence of ossification appears relatively similar to that observed in other marsupials (Smith, 1997; Sánchez-Villagra et al., 2008). The more advanced state of *M. domestica* at birth, compared with the dunnart, can largely be explained by a somewhat longer gestation length in the opossum; about 1.5 to 2 days longer than the smaller dunnart (Cook et al., 2021). As with other marsupials, immediately following birth, several organ systems become crucial to neonate survival such as the lungs, integument, digestive tract, and excretory system (Smith and Keyte, 2020). Although many of these systems are rudimentary at birth in *M. domestica* (Fig. 2A), they are comparatively better developed than in other marsupials. For instance, although the eye is poorly developed at birth in *M. domestica*, distinct retinal pigmentation is present, unlike in the dunnart (Cook et al. 2021). Additionally, whereas the highly translucent integument in the dunnart, indicative of cutaneous gas exchange, remains until at least postnatal day 10 (Cook et al., 2021), in *M. domestica*, skin rapidly becomes opaque after birth and by P5 the lungs exhibited increased vascularisation suggesting greater reliance on pulmonary gas-exchange. From birth until weaning, other postnatal events show similar evidence of accelerated initiation. Hindlimb and tail separation (P5), digit separation, mouth delamination, and body fur establishment, all occur more than 5 days in advance of events in the dunnart. Acceleration and earlier initiation is especially apparent in traits linked to thermoregulation such as body size and pelage formation. Although the general trajectory of body growth is similar between *M. domestica* and the dunnart, the rate of body elongation appears distinctly more rapid from P10 onwards in *M. domestica*. In *M. domestica*, hair appears on the head at P7, and later, on the rump from P15 (Fig. 1). Although hair first appears on the dunnart head at a similar postnatal time (P8), body hair does not appear until five days later (P20). Other characters also linked to precociality also initiate earlier. These include sensory systems like the eyes, ears, and vibrissae, but also locomotor traits such as grasping features of the hands and feet, and tail mobility. For instance, the eyes of *M. domestica* open at least 30 days earlier, and the separation and unfolding of the ear pinnae, starts about 15 days earlier than in the dunnart. In the postcranial region, whereas the hindlimb and tail are still fused in the dunnart until P9, in *M. domestica* these structures are distinct by P6. General rate of ossification in the hindlimb is also more advanced, with tarsal ossification at P40 in *M. domestica* (Fig. 2B) being comparable to that of the dunnart at P50 (Cook et al., 2021).

Overall, our observations in *M. domestica*, and comparison with marsupials possessing a pouch suggest that rates of postnatal development are generally accelerated, or occur earlier, when the pouch is absent (Fig. 2B). Accelerated and earlier initiating precocial traits probably serve at least two main functions. First, it unburdens the mother earlier from the costs of rearing and carrying offspring. Second, it increases survival of the offspring by reducing the risks of detachment, predation and thermal shock that are heightened without the pouch present. Comparison of variation life history and reproductive traits across marsupial species (Fig. 3) reveals a pronounced gradient connecting more r-selected, pouchless marsupials, possessing relatively short weaning intervals, large litter sizes and accelerated precociality (e.g., detachment, eye opening age) such as many didelphids, with more K-selected, pouched marsupials that produce only single (rarely two) offspring such as kangaroos and vombatiforms. Dasyuromorphians, which exhibit a range of pouch structures (Edwards and Deakin, 2012), are intermediaries with large to modest sized litters, but also relatively altricial offspring (Tyndale-Biscoe, 2005; Cook et al. 2021). This life history gradient also broadly correlates with body size, with all K-selected, pouched taxa being members of clades with large-bodied representatives, whereas r-selected strategies are dominated by small sized ameridelphian and dasyuromorphian marsupials.

**Figure 3.**
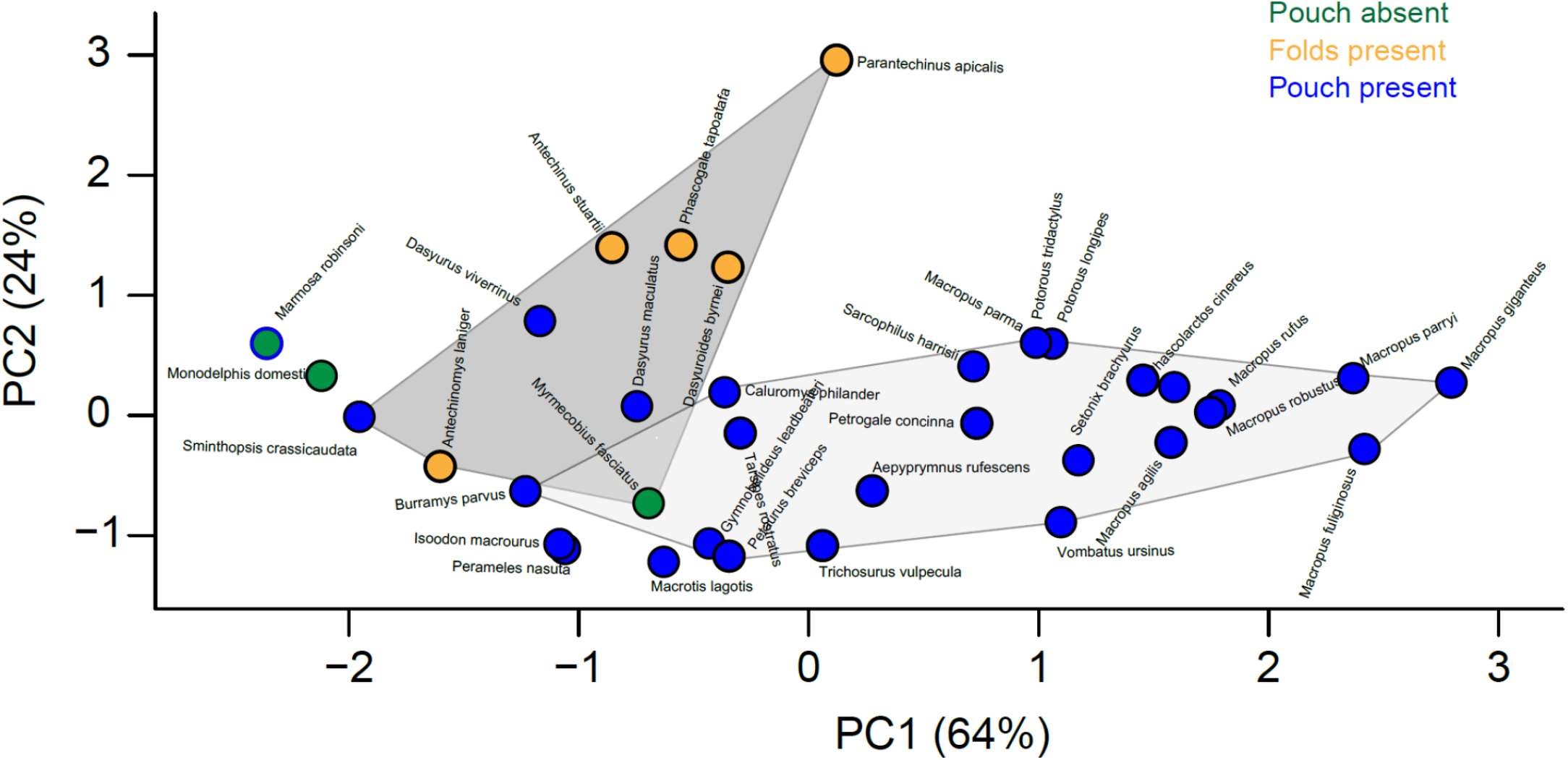
Principal component analysis of life history and reproductive traits across 36 marsupial species. Dark grey hull = Dasyuromorphia. Light grey hull = Diprotodontia.

The relative altriciality of marsupial young is their most striking difference from their sister radiation, the placental mammals (Tyndale-Biscoe and Renfree, 1987). Traditionally, this mode of development has been viewed as potentially intermediate between the egg-laying monotremes and extensive gestation of placentals (Lillegraven 1969). However, increasing evidence suggests that marsupial altriciality is derived amongst marsupials (White et al., 2023). This is based both on the recognition of the highly specialised adaptations needed to support neonatal survival at what would otherwise be embryonic stages of mammalian development (Smith and Keyte, 2020), as well as fossil evidence that non-metatherian mammals like multituberculates and placentals had growth rates more characteristic of precocial life histories (Weaver et al., 2022). Phylogenetic results here support the highly derived nature of marsupial reproduction and life history. We recover evidence across the best-ranked character models that the pouch evolved within crown-group marsupials, as opposed to more deeply in mammalian tree, and that the pouch as evolved in parallel at least twice (Fig. 4). Specifically, the pouch as found to be absent amongst the ancestor of crown-group marsupials, with separate acquisitions amongst Didelphimorphia and Australidelphia. The pouch or pouch-like structures as homoplastic is consistent with the large structural variation in pouch morphology across marsupials (Edwards and Deakin, 2012). Our results thus support Kirsch’s (1977) argument that, despite its familiar association with marsupials, the pouch is not ancestral to the clade. Furthermore, because of the close link we identify between the presence of a pouch, and reproductive and life history traits, the inference of pouch absence amongst ancestral marsupials also suggests a much more r-selected life history, perhaps extending also to more basal metatherians.

**Figure 4.**
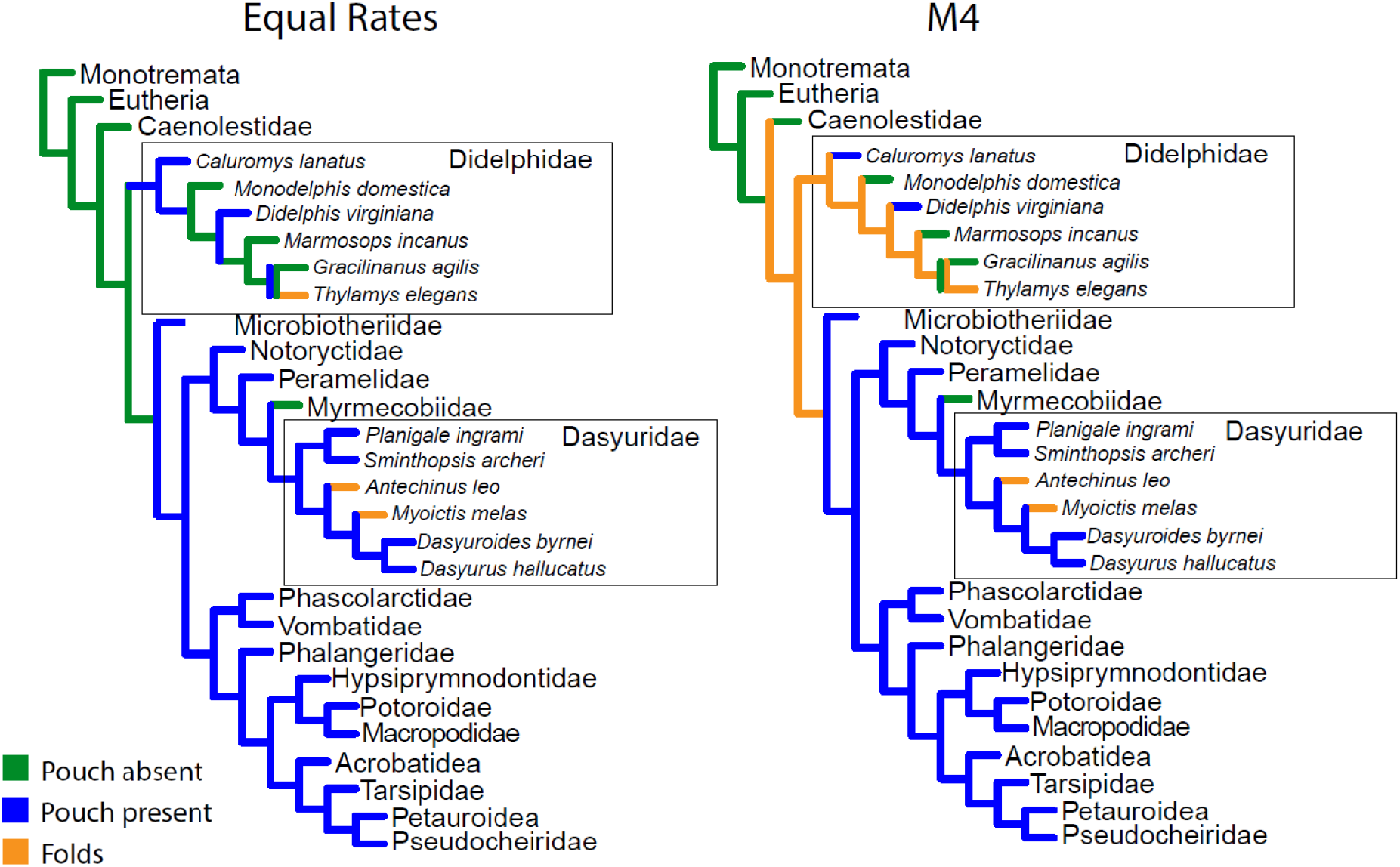
Family level phylogenetic consensus (>50% maps) of the best ranked models of pouch evolution based on 500 stochastic character maps each on 100 randomly sampled trees. An equal rates model recovers the pouch as absent amongst crown-group marsupials as well as amongst the last common ancestor of Didelphimorphia and Australidelphia. Under an M4 character transition model, marsupials ancestrally possess abdominal folds, from which true pouches were derived seperately in Austalidelphia and Didelphimorphia.

The parallel origins of a pouch, and by inference reproductive and life history strategies amongst australidelphian and ‘ameridelphian’ marsupials is also congruent with the biogeographic partitioning of marsupial clades during a phase when Australia and South America were geographically isolated (Lagabrielle et al., 2009; Crespo and Goin 2025). Amongst didelphimorphians, the phylogenetic reconstructions suggest a pouch had evolved amongst some groups by at least the Early Miocene, and perhaps the Late Eocene. Amongst australidelphians, the stochastic character mapping suggests that pouch origins extend to the clades inception, perhaps the Early Palaeocene. The most basal australidelphian, the tiny South American microbiotherian, *Dromiciops australis*, possesses a pouch, and generally gives birth to no more than four, and typically three, offspring (Nowak, 1999). Whether the reproductive strategy of *D. australis* is reflective of the ancestral australidelphian condition is unclear. The consensus phylogenetic reconstruction suggests that early australidelphians may have lacked a pouch or only had an incipient one (Fig. 5), presumably possessing a more r-selected reproductive strategy akin to an ancestral marsupial. If Australidelphia originated within South America, before dispersal to Antarctica and Australia, as some biogeographic scenarios propose (Nilsson et al., 2010), such a strategy may have been favourable in traversing the archipelago hypothesised to link west and east Gondwana during the Late Cretaceous and early Paleogene (Crespo and Goin, 2025).

**Figure 5.**
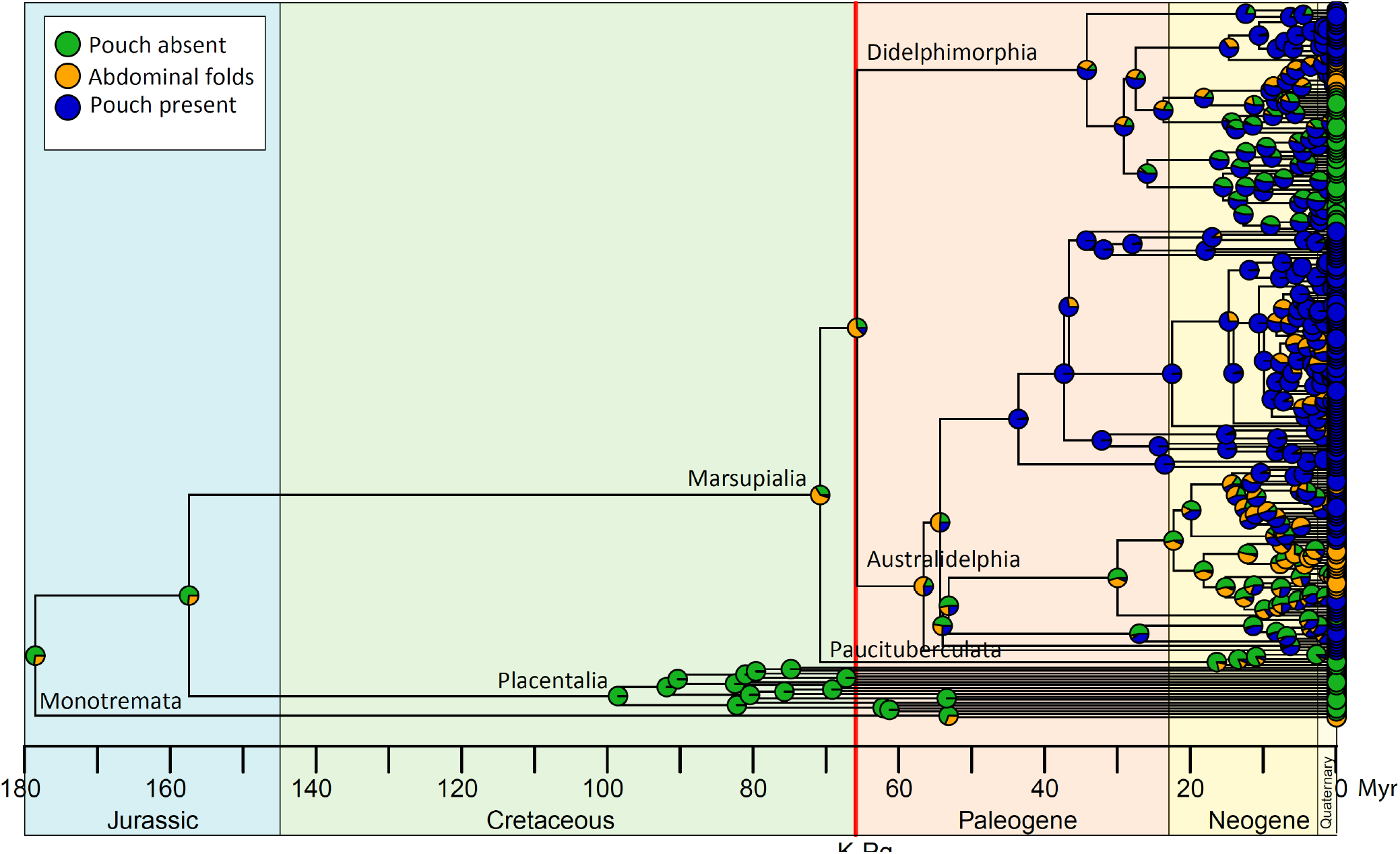
Ancestral state reconstruction of pouch states on a timescaled phylogeny of marsupials and outgroup mammalian taxa.Summary of 500 stochastic character maps of pouch states under an ‘M4’ character model across 100 phylogenetic trees (244 extant mammalian taxa, with 226 marsupial species). Timescale is millions of years before present (Myr).

Ancestral state reconstructions based on the best ranked models of character evolution recover a high rate of character transitions from a ‘non-pouch’ to ‘pouch’ state near the Cretaceous–Paleogene boundary (Fig. 6). Summary analysis recovers a split between Didelphimorphia and Australidelphia in the immediate aftermath of the K–Pg, with two basal marsupial branches crossing the boundary without a pouch or possessing some intermediate state (Fig. 5). A post-K–Pg radiation of at least Australidelphia, and possibly all living marsupials has been recovered by several phylogenetic analyses (e.g., Nilsson et al., 2003; Mitchell et al., 2014). Aside from the most basal crown-group marsupial lineages which may have crossed the boundary, fossil evidence suggests that basal metatherians suffered a high rate of extinction compared with eutherian and multituberculate lineages; as high as 90% (Alroy, 1999; Lillegraven and Eberle, 1999; Wilson, 2013). Lillegraven et al. (1987) hypothesised that marsupials may have been more severely impacted due to: (1) increased sensitivity of their offspring to environmental fluctuations; and (2) reduced adaptability or competitive inferiority relative to eutherians. Environmental disturbances associated with an asteroid impact at the boundary are thought to have been severe, including potentially a brief thermal pulse (Robertson et al., 2004), followed by an extended period of global cooling and darkness(Chiarenza et al. 2020). Results here are not consistent with the hypothesis that marsupial declines are explicable due to vulnerability to environmental change. Rather, our data suggest that boundary crossing basal marsupial lineages, and perhaps some basal metatherians, likely possessed life history traits (i.e., large litters, short gestation and weaning intervals, fast sexual maturity) which should have increased robustness to environmental disruptions of the impact, and indeed may have been a key factor in their persistence across the boundary. Lillegraven et al’s second hypothesis is more compelling. Marsupials with r-selected life history strategies are smaller bodied and less ecologically and anatomically disparate than marsupials with a pouch. Based on comparison of pouchless *M. domestica* with pouched australidelphian taxa (Fig. 2B), we suggest that this discrepancy could be largely explained by the longer postnatal development available to marsupial possessing a pouch. This extended postnatal rearing allows for extended growth, a key requirement for many character innovations (i.e., elongate limbs, digits, larger size), especially in later forming more caudal regions of the body. Marsupials with the most extensive caudal innovations, such as macropodids, also have the most elongated postnatal attachment, weaning, and sexual maturity intervals (Fig. 2B), and the characters which appear latest in their ontogeny (e.g., molars, hind-limb, tail) are also the most derived relative to marsupials with shorter postnatal periods. Thus, although an r-selected life history strategy may have been advantageous for metatherian persistence through the end-Cretaceous extinction, it may also have limited their ability to acquire key caudal innovations needed to fully exploit the ecological vacancy of the post-extinction world.

**Figure 6.**
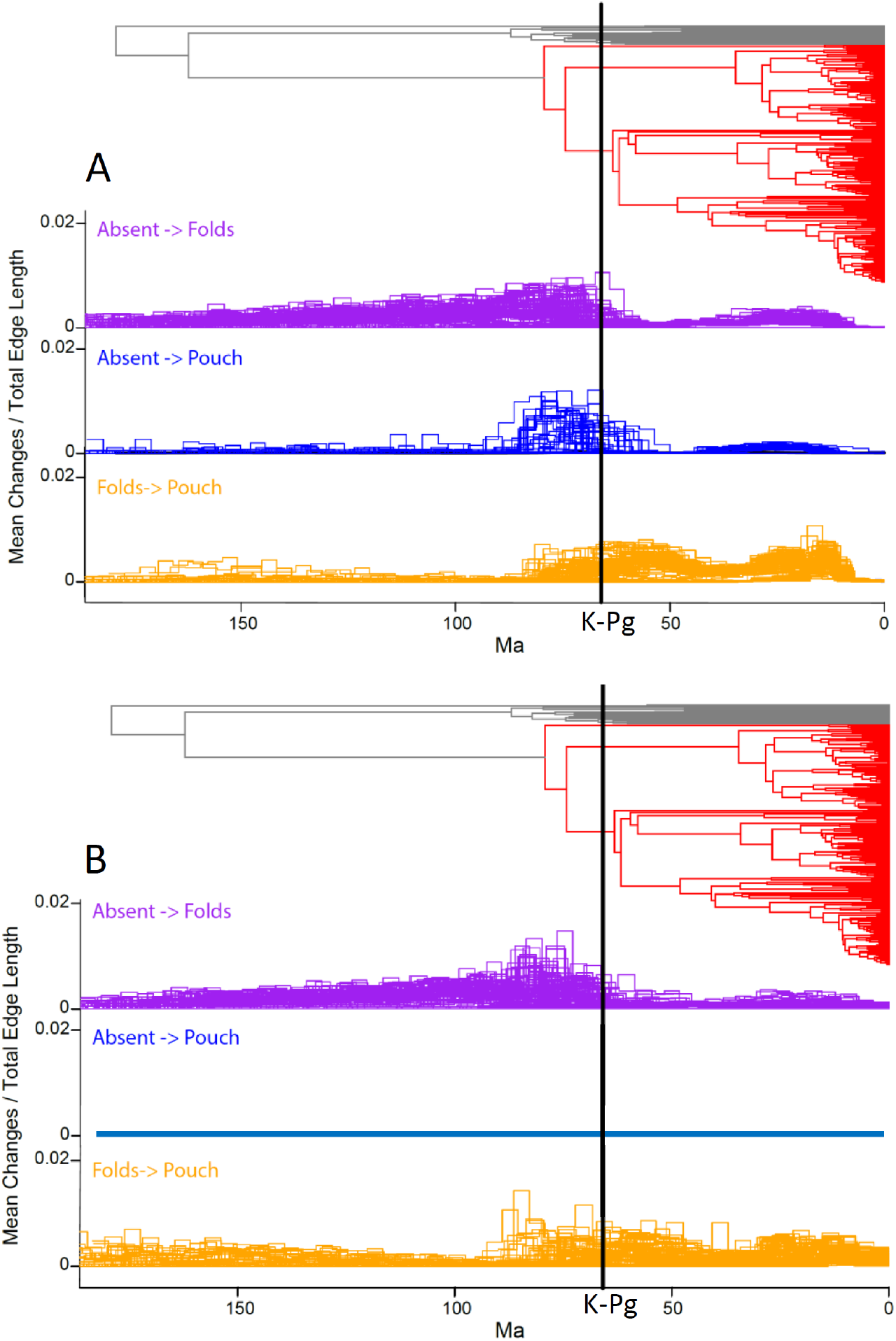
Rates of character state change through geological time (millions of years before present, Ma) under two best ranked transition models. **A.**Equal rates transition model. **B**. M4 rates transition model. Summary of 500 stochastic maps each on 100 randomly sampled trees. Consensus tree topology (marsupials in red) from Upham et al. (2019).

## Acknowledgements

For scanning we thank Tautis Skorka at the University of Southern California. Assistance with questions about ruthenium red staining methodology was kindly provided by Stephan Handschuh. We thank staff from the Division of Laboratory Animal Medicine (UCLA) for assisting with animal care and veterinary support. Sears laboratory members are thanked for feedback on aspects of this work. J. Marcot is kindly acknowledged for providing financial support for AMCC during his postdoctoral research. AMCC also gratefully acknowledges financial support provided by an American Association for Anatomy Postdoctoral Fellowship. This work was supported by grants from the National Science Foundation and National Institute of Health to K. Sears.

